# Empirical Performance of Tree-based Inference of Phylogenetic Networks

**DOI:** 10.1101/693986

**Authors:** Zhen Cao, Luay Nakhleh

**Affiliations:** Department of Computer Science, Rice University

## Abstract

Phylogenetic networks extend the phylogenetic tree structure and allow for modeling vertical and horizontal evolution in a single framework. Statistical inference of phylogenetic networks is prohibitive and currently limited to small networks. An approach that could significantly improve phylogenetic network space exploration is based on first inferring an evolutionary tree of the species under consideration, and then augmenting the tree into a network by adding a set of “horizontal” edges to better fit the data.

In this paper, we study the performance of such an approach on networks generated under a birth-hybridization model and explore its feasibility as an alternative to approaches that search the phylogenetic network space directly (without relying on a fixed underlying tree). We find that the concatenation method does poorly at obtaining a “backbone” tree that could be augmented into the correct network, whereas the popular species tree inference method ASTRAL does significantly better at such a task. We then evaluated the tree-to-network augmentation phase under the minimizing deep coalescence and pseudo-likelihood criteria. We find that even though this is a much faster approach than the direct search of the network space, the accuracy is much poorer, even when the backbone tree is a good starting tree.

Our results show that tree-based inference of phylogenetic networks could yield very poor results. As exploration of the network space directly in search of maximum likelihood estimates or a representative sample of the posterior is very expensive, significant improvements to the computational complexity of phylogenetic network inference are imperative if analyses of large data sets are to be performed. We show that a recently developed divide-and-conquer approach significantly outperforms tree-based inference in terms of accuracy, albeit still at a higher computational cost.

## 1 Introduction

As evidence of reticulation (hybridization, horizontal gene transfer, etc.) in the evolutionary histories of diverse sets of species across the Tree (or, more appropriately in our context, Network) of Life continues to grow, increasingly sophisticated methods for phylogenetic network inference are being developed to incorporate processes such as incomplete lineage sorting (ILS), gene duplication and loss, and gene flow [20, 21, 15, 16, 24, 28, 27]. These methods, which are mostly statistical in nature, are computationally prohibitive due, in part, to the complex space of phylogenetic networks that is explored. This, in turn, has limited the applicability of such methods to data sets with small numbers of taxa, loci, and reticulation events. To ameliorate this computational challenge, a divide-and-conquer approach was recently introduced [26], where this accurate, yet computationally expensive, method could be used to infer small networks that are then merged to produce a phylogenetic network on the full data set. The accuracy of the method notwithstanding, the inference of small networks remained a computational bottleneck.

An alternative approach that could be considered would first infer an underlying “species tree”, and then augment this tree into a network by adding reticulations to it to fit the data under some criterion that incorporates reticulation. The benefit of such an approach is that it would utilize one of a wide array of species tree inference methods that are both accurate and efficient, thus drastically reducing the space of phylogenetic networks to explore by limiting them to those “based” on inferred trees. Indeed, the question of whether a species tree can be accurately inferred in the presence of reticulation has been partially explored from theoretical [12] as well as empirical [4] angles. An extensive simulation study on small data sets demonstrated the problems with inferring “the” species tree in the presence of high rates of gene flow [13]. However, a more general question to ask is whether tree inference methods can infer a tree, not necessarily the species tree (if one insists on using this designation), that can be augmented into the correct network. Along the same lines, the class of tree-based networks was introduced [6] and the set of trees that characterize a phylogenetic network in the presence of ILS was revisited [29].

In this paper, we study the performance of tree-based inference of phylogenetic networks by exploring two popular methods for inferring a start tree and two network criteria that scale to evaluating large networks. We consider synthetic networks that are generated under a birth-hybridization model and data simulated under the multispecies network coalescent [19]. We find that the concatenation method does poorly at inferring a backbone tree of the network, whereas the commonly used method ASTRAL [23] performs significantly better at this task. However, even when a correct network-backbone tree is used, augmenting such a tree into the correct network is a challenging task and results in poor accuracy under both the pseudo-likelihood and minimizing deep coalescences criteria of [21, 22]. It is worth noting that the size of phylogenetic networks we consider in our study makes use of the likelihood criterion of [20] infeasible. We demonstrate that the divide-and-conquer approach of [26] yields more accurate results, yet is orders of magnitude slower than tree-based inference. We demonstrate that combining the strengths of the two—the speed of tree-based inference and the accuracy of the divide-and-conquer approach—could provide a promising approach to large-scale network inference. Finally, our implementations of tree-based phylogenetic network inference are implemented in PhyloNet [14, 17].

## 2 Background

A **phylogenetic network** Ψ on a set 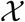 of taxa is a rooted, directed, acyclic graph (DAG) whose leaves are bijectively labeled by 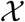. For Ψ = (*V*(Ψ), *E*(Ψ)), the set *V*(Ψ) of nodes contains a root node with in-degree 0 and out-degree 2, tree nodes with in-degree 1 and out-degree 2, reticulation nodes with in-degree 2 and out-degree 1, and leaf nodes with in-degree 1 and out-degree 0. If *v* is a reticulation node, then the two edges incident into it are reticulation edges, and all edges incident into tree nodes are tree edges. In a statistical setting, reticulation edges are associated with inheritance probabilities, and all edges have lengths, so that the phylogenetic network defines a distribution over gene trees under the multispecies network coalescent [20].

A **phylogenetic tree** *T* can be viewed as a phylogenetic network with no reticulation nodes. There is a natural relationship between a phylogenetic network and the set of trees it displays.

### Definition 1

A tree T on set 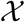 of taxa is displayed by a phylogenetic network Ψ (also on set 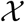 of taxa) if T can be obtained from Ψ by removing a set of reticulation edges followed by forced contraction of every node v of in- and out-degree 1, where the edges (u, v) and (v, w) are replaced by a single edge (u, w) and v is deleted.

We denote by 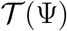 the set of all trees displayed by Ψ.

Several methods for inference of phylogenetic networks in the presence of incomplete lineage sorting have been devised. These include parsimony methods [18], maximum likelihood methods [19, 20], maximum pseudo-likelihood methods [21, 27], and Bayesian methods [15, 16, 24, 28]. These methods have poor scalability in terms of time and memory requirements and are applicable to very small data sets. The pseudo-likelihood methods were devised to ameliorate the problem of computing the full likelihood of phylogenetic networks, but these suffer from the large network space they need to explore.

One potential approach to tackling the computational requirements of phylogenetic network inference is based on first inferring a “species tree” and then adding a set of reticulation edges to it, in the hope of obtaining the true network. Indeed, as discussed above, the plausibility of this approach has been addressed, albeit in a limited fashion [4, 12, 6, 29, 13]. If this approach works in practice, it would tremendously reduce the phylogenetic network space to search and, consequently, scale phylogenetic network inference to much larger data sets. The goal of this work is to systematically study this approach on phylogenetic networks generated under a birth-hybridization model to assess its potential. Next, we describe our implementation of this approach.

## 3 Methods

Consider a data set 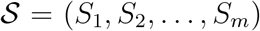 and 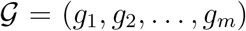, where *S_i_* is the alignment of a set of orthologous sequences of locus *i* in the genomes of a set 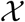 of taxa, and *g_i_* is a rooted gene tree for locus *i* inferred from *S_i_*. Throughout this work we assume that loci are independent and each locus is recombination-free.

Generally, a backbone-based approach to inferring a phylogenetic network Ψ on set 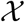 of taxa follows two steps.

**Step 1.** Build a start tree *T*, either using the sequences 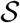 directly or using the estimated gene trees 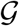.
**Step 2.** Augment the start tree *T* into a network Ψ by adding a set *E_r_* of reticulation edges to it to optimize some criterion given either 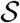 or 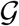.

A desired property of the start tree *T* in Step 1 is that it is a backbone of the network, that is, it is displayed by the true network so that the latter is obtainable by the backbone-based approach. This property is illustrated in Figure 1.

**Figure 1:**
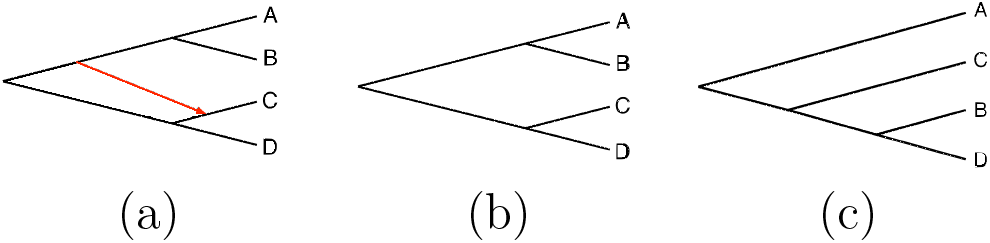
From a backbone tree to a phylogenetic network. (a) Phylogenetic network Ψ. (b) A backbone tree that is displayed by Ψ and, thus, can be augmented into the true network. (c) A tree that cannot be augmented to produce Ψ and therefore is not a backbone of the network.

### 3.1 Step 1: Building a start tree

As we only consider reticulation and incomplete lineage sorting (ILS) in this paper (that is, we do not consider gene duplication and loss, for example), we considered two methods that are heavily used to infer a species tree when incongruence is suspected to be due to ILS. The first method infers a tree on the concatenation of the sequence data 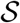. For this purpose, we used IQ-TREE [9] to infer a tree from the concatenated sequences. The second method infers a species tree from individual gene trees in 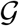. For this purpose, we used ASTRAL-III [23]. Both of these methods have been shown to have good efficiency and accuracy in practice.

### 3.2 Step 2: Augmenting the start tree into a network

Let Ψ be a phylogenetic network (including the case where Ψ is a tree). Adding a reticulation edge to Ψ is done by selecting two edges (*u, v*) and (*x, y*) in Ψ, replacing them by (*u, m*), (*m, v*), (*x, m′*), and (*m′, y*), and adding an edge (*m, m′*). This operation results in creating a new reticulation node, *m′*, in the phylogenetic network Ψ and results in a network with one more reticulation node than those in Ψ. The reticulation edge can be added to initial edges or added edges. Hence, obtained networks are not limited to tree-based networks.

Consider an optimality criterion Φ that is defined on a given network Ψ and input data 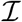 (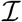 could be sequences or trees, and the criterion has to be appropriate for the type of data used). Given a maximum number of reticulation nodes to consider *MAX*, one implementation of tree-based inference of networks follows the steps of Algorithm **StepwiseAugment**, given a start tree *T*.

#### Algorithm 1: StepwiseAugment

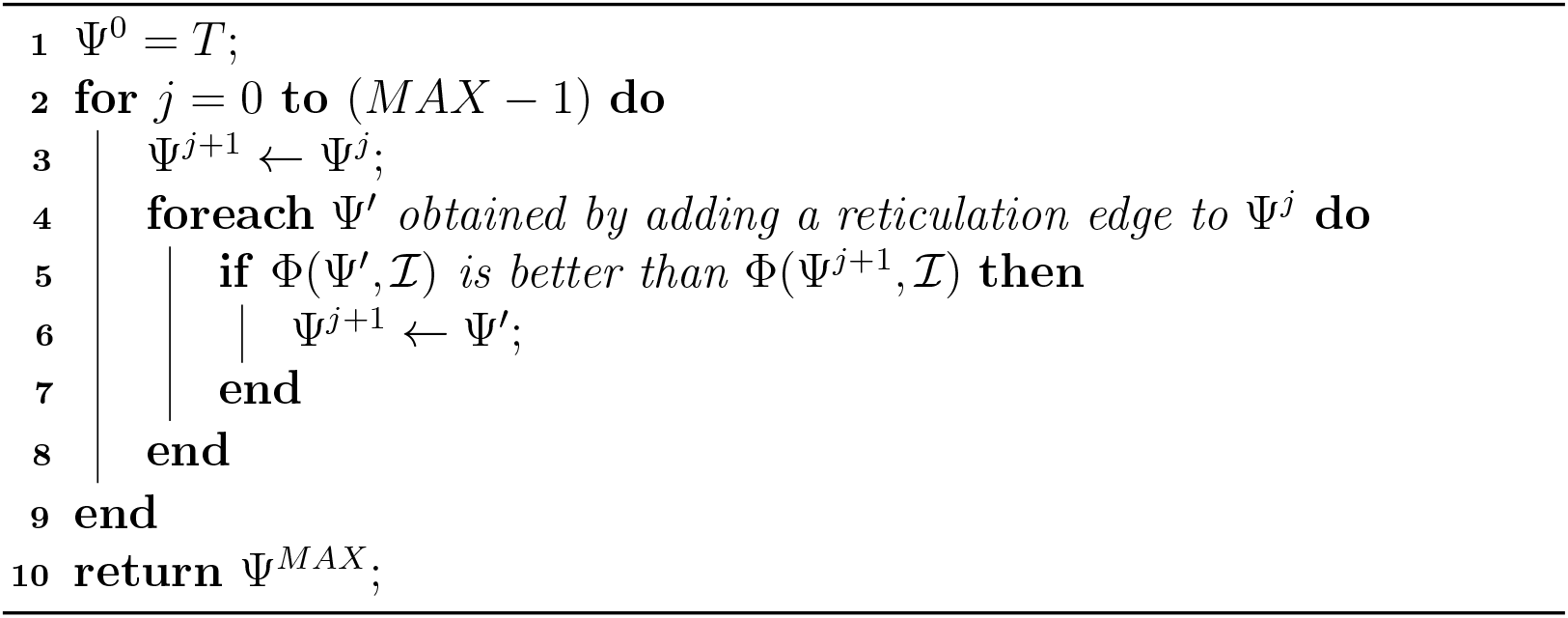

In other words, we exhaustively evaluate the optimal networks with *j* + 1 reticulations that can be obtained by adding a single reticulation to the optimal network found with *j* reticulations. It is important to highlight here that this implementation is not the most exhaustive way to augment a tree into a network with a given number of reticulations. The most exhaustive way of augmenting a tree on *n* leaves into a phylogenetic network with *k* reticulations considers 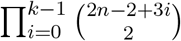 networks, which is *O*(*n*^2*k*^) when *k* ≪ *n*. Our implementation reduces this number to 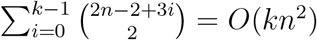.

Another approach to augment a start tree into a network is to heuristically search for reticulation edges to add in a local search fashion. We implemented **LocalSearchAugment** which starts with a tree *T* and then randomly chooses one of a set of operations to apply in order to augment the current network into a new candidate network. If the resulting network has a better optimality score, it is adopted as the new current network and the search continues; otherwise, the move is rejected (or rejected with a probability) and a new candidate network is considered. The set of moves we considered consist of: (i) adding a new reticulation edge; (ii) removing one of the reticulation edges that have been added; (iii) relocating the head of an added reticulation edge; (iv) relocating the tail of an added reticulation edge; (v) reversing the direction of an added reticulation edge; (vi) replacing an added reticulation edge with a new one; and, (vii) modifying an edge parameter like the inheritance probability and branch length in the network (if criterion Φ requires such parameters). We define a probability distribution on this set of moves, and in each iteration of the local search, a move is selected randomly based on this distribution. The heuristic search applies random restarts to ameliorate the local optima issue, where each search stops when the number of consecutively rejected moves (they would be rejected because the proposed solution candidates do not improve the optimality criterion) reaches a pre-set threshold. It is important to emphasize that **LocalSearchAugment** never removes any of the edges that exist in the start tree *T*.

For Φ, we considered two criteria in this study: the pseudo-likelihood criterion of [21] and the minimizing deep coalescence (MDC) criterion of [18]. Both of these criteria use gene trees as input data. The likelihood criterion of [20] is infeasible to compute on the data sets we consider in this study.

### 3.3 Evaluating the inferred trees and networks

While several dissimilarity measures exist for comparing networks and, by extension, trees and networks, e.g., [2, 3, 8], these measures are very sensitive to the misplacement of reticulation nodes, even in cases when networks agree on much of their structure. Therefore, in this paper, we used two measures of dissimilarity for tree-to-network and network-to-network comparisons.

Letting RF be the Robinson-Foulds distance [11] (the size of the symmetric difference between the two trees, divided by 2), the dissimilarity between a tree *T* and a network Ψ is:

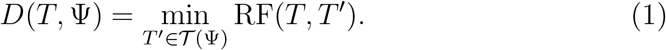

For network-to-network comparison, we can extend the notion of displaying to networks: We say phylogenetic network Ψ′ is displayed by phylogenetic network Ψ if Ψ′ can be obtained by removing a set of reticulation nodes and applying forced contraction to Ψ (but unlike in the case of trees, some reticulation nodes remain in the network). We denote by 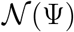 the set of all networks displayed by Ψ (obviously, 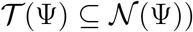. We denote by *r*(Ψ) the number of reticulations and 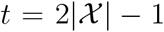 the number of nodes for any displayed tree of Ψ. Let Ψ_*t*_ and Ψ_*i*_ be true and inferred networks, respectively, and 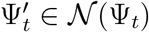 and 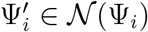 be the two displayed networks that have the smallest distance 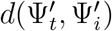, as computed by [8], over all displayed networks of the two networks Ψ_*t*_ and Ψ_*i*_, and 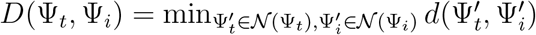. If more than one pair of displayed networks have the smallest distance, a pair with the largest number of reticulation nodes is selected. We now define:

- True positives: 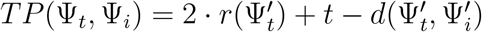.
- True positives rate: *TPR*(Ψ_*t*_, Ψ_*i*_) = *TP*(Ψ_*t*_, Ψ_*i*_)/(*t* + 2 · *r*(Ψ_*t*_)).
- False positives: 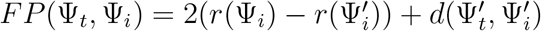.
- False positives rate: *FPR*(Ψ_*t*_, Ψ_*i*_) = *FP*(Ψ_*t*_, Ψ_*i*_)/(*t* + 2 · *r*(Ψ_*i*_)).
- False negatives rate: *FNR*(Ψ_*t*_, Ψ_*i*_) = 1 − *TPR*(Ψ_*t*_, Ψ_*i*_).

For example, if the networks of Figure 1(a) and Figure 1(c) are the true network Ψ_*t*_ and inferred network Ψ_*i*_, respectively, then 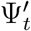 and 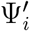 are the networks of Figure 1(b) and Figure 1(c), respectively. In this case, we have 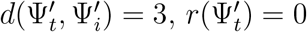, *TP* = 4, *FP* = 3, *TPR* = 4/9, and *FPR* = 3/7.

## 4 Results and Discussion

To assess the performance of tree-based inference of phylogenetic networks, we tested both methods described above on simulated data. Furthermore, we analyzed a biological data set and contrasted the results to those obtained by other methods.

For the simulated data, we used the same 24 networks used in [26], as those were generated under a birth-hybridization model and varied in their complexity. For each integer in [0, 5], four networks in the data set have that number of reticulation nodes. Two of the 24 networks are not tree-based [25].

As in [26], these networks were divided into 3 groups of hardness (in terms of inference): 8 “easy” (E), 8 “medium” (M), and 8 “hard” (H). Each network has 16 taxa and 1 outgroup. For each network, we generated 100 gene trees with two individuals per species using the program `ms` [7], and then generated sequences of length 1000 using Seq-gen [10] under the GTR model. We set the population mutation rate at 0.02, base frequencies of A, C, G and T at [0.2112,0.2888,0.2896,0.2104], respectively, and the transition probabilities at [0.2173,0.9798,0.2575,0.1038,1,0.2070]. We then ran the aforementioned methods on the data. In terms of the complexity of the data sets, the number of distinct gene trees (out of 100) in the 24 data sets varied between 69 and 100. That is, the rate of ILS is quite high.

### 4.1 Accuracy of the start tree

#### 4.1.1 Performance of the concatenation method

By concatenating the sequence data of all loci, we obtained for each network 34 sequences of length 100000 each (since we have two individuals per species). We then inferred a tree on the concatenated sequences of each of the 24 data sets using IQ-TREE [9]. As there are 34 taxa in the resulting trees but there are only 17 species, we first identified the data sets where the two individuals from each of the 17 species form a monophyletic group. We found that for 21 data sets, the resulting tree group the two individuals from each species monophyletically, whereas in the three remaining data sets, the individuals of some species were not monophyletic. For the 21 data sets, we “collapsed” the two individuals in the inferred tree into a single leaf with the respective taxon name so as to compare the accuracy of the inferred “species tree” to the true phylogenetic network.

Of the 21 data sets, we found that 12 of them resulted in species trees that are displayed by, or are backbone trees of, their corresponding true network. The remaining nine trees had an average distance, based on Eq. (1), of 3.33.

These results illustrate that in the presence of ILS and hybridization, concatenation has a poor performance; only in half of the 24 data sets did the method infer a tree that could be augmented into the true network.

#### 4.1.2 Performance of ASTRAL

We then turned to assessing the accuracy of species trees inferred by ASTRAL-III [23]. We inferred a gene tree on the sequence alignment of each locus using IQ-TREE [9] and used these inferred gene trees as the input set 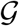 of gene trees to ASTRAL-III.

ASTRAL-III had a much better performance than species tree inference on the concatenated sequences. In fact, ASTRAL-III inferred a backbone tree that could be augmented into the respective true network in 87.5% of the data sets. More specifically, 21 out of 24 species trees inferred by ASTRAL-III are backbone trees of the true networks. For the 3 data sets where the inferred tree could not serve as a backbone of the true network, the average distance between the trees and the true networks, as computed by Eq. (1), was 2.67.

This shows that the start tree built from inferred gene trees using ASTRAL-III is much better than concatenation using IQ-TREE. This is because the rate of ILS is high, and ASTRAL-III considers the gene tree topology conflicts.

To explore whether the performance of ASTRAL-III would improve with more loci, we increased the number of gene trees simulated on each network to 1,000 and used the true gene trees as input to ASTRAL-III. Now, ASTRAL-III inferred a tree that is the backbone of its respective true network for 22 of the 24 data sets.

### 4.2 Accuracy of the inferred networks

Given the accuracy of the species trees produced by ASTRAL-III, we used it as the start tree for network inference. Since the input gene trees and output species trees of ASTRAL-III are all unrooted, we rerooted all the trees at the designated outgroup and deleted it. Then we used the rooted gene trees and rooted species trees as input to both network inference procedures. We evaluated the performance of the network inference on NOTS (Night Owls Time-Sharing Service), which is a batch scheduled High-Throughput Computing (HTC) cluster.

#### 4.2.1 Performance of StepwiseAugment

We ran **StepwiseAugment** on the start trees obtained by ASTRAL-III on all 24 data sets, under the MDC criterion, while specifying to the inference method the true number of reticulations. While knowing the true number of reticulations is not doable in practice, we made this decision for two reasons. First, we wanted to test how the approach works under the ideal situation (of knowing the true number of reticulations). Second, determining the number of reticulations is not doable in a systematic way with the MDC criterion. The correctness achieved by **StepwiseAugment** in this case is 41.7%, reflecting that in only 10 out of the 24 cases did the method infer the correct networks. For the 3 data sets where the start trees are not backbone trees of the networks, obviously, the method could not infer the correct network. For the 17 data sets with at least one reticulation and correct start trees, the method was able to find the correct reticulation in only 8 of them. These results show that **StepwiseAugment** coupled with the MDC criterion is not a viable approach for inferring phylogenetic networks.

#### 4.2.2 Performance of LocalSearchAugment

We then set out to study the performance of **LocalSearchAugment** when using the ASTRAL-III species tree as the start tree and under both the pseudo-likelihood and MDC criteria. One might ask: If the “quasi brute-force” procedure **StepwiseAugment** did not perform well, could a local search heuristic perform better? The answer in our case is positive. The reason for this is because when searching for an optimal network with *k* + 1 reticulations, **StepwiseAugment** is already “stuck” with the *k* reticulations it had identified already, and this might not necessarily result in an optimal solution. A truly brute-force approach for considering all networks with *MAX* reticulations on a start tree *T* would first identify the set of all possible reticulations that could be added to *T* (including “dependent” reticulations, where a reticulation edge is added between two reticulation edges or between one tree and one reticulation edge), and then consider every subset of them. Such an approach is not only computationally infeasible for the size of the data sets we consider, but is also very hard to implement because of the dependent reticulations.

Since inference under neither MDC nor pseudo-likelihood is equipped with a systematic way to determine the number of reticulations (i.e., a stopping rule), we ran **LocalSearchAugment** under both criteria in two different ways. In one way, we set the number of reticulations for each data set to be the true number for that data set. In the figures below, we label results based on this setting as MPL (for maximum pseudo-likelihood) and MDC (for minimizing deep coalescences). In the other way, we set the maximum number of reticulations at 5, which is the largest number of reticulations in all 24 data sets. In the figures below, we label results based on this setting as MPL’ (for maximum pseudo-likelihood) and MDC’ (for minimizing deep coalescences). The local heuristic search on each data set was repeated 20 times with random restarts to obtain the network estimates.

We also ran MPL on 1,000 gene trees to explore its convergence when the amount of data is larger. In the figures below, results of this run are labeled MPL1000. Finally, we compared these methods to two methods that can scale to larger data sets: The maximum pseudo-likelihood method of [21], where the network space is searched directly without a fixed start tree (the maximum number of reticulations is set at the true value), and the newly developed divide-and-conquer method of [26] when the full set of trinets is considered in the divide step (this method does not require a pre-set number of reticulations). The results of these runs are labeled MPLo and *D*&*C*, respectively, in the figures below. For these two methods, the input gene trees is the set of gene trees inferred by IQ-TREE.

The results of all these runs are summarized in Figure 2.

**Figure 2:**
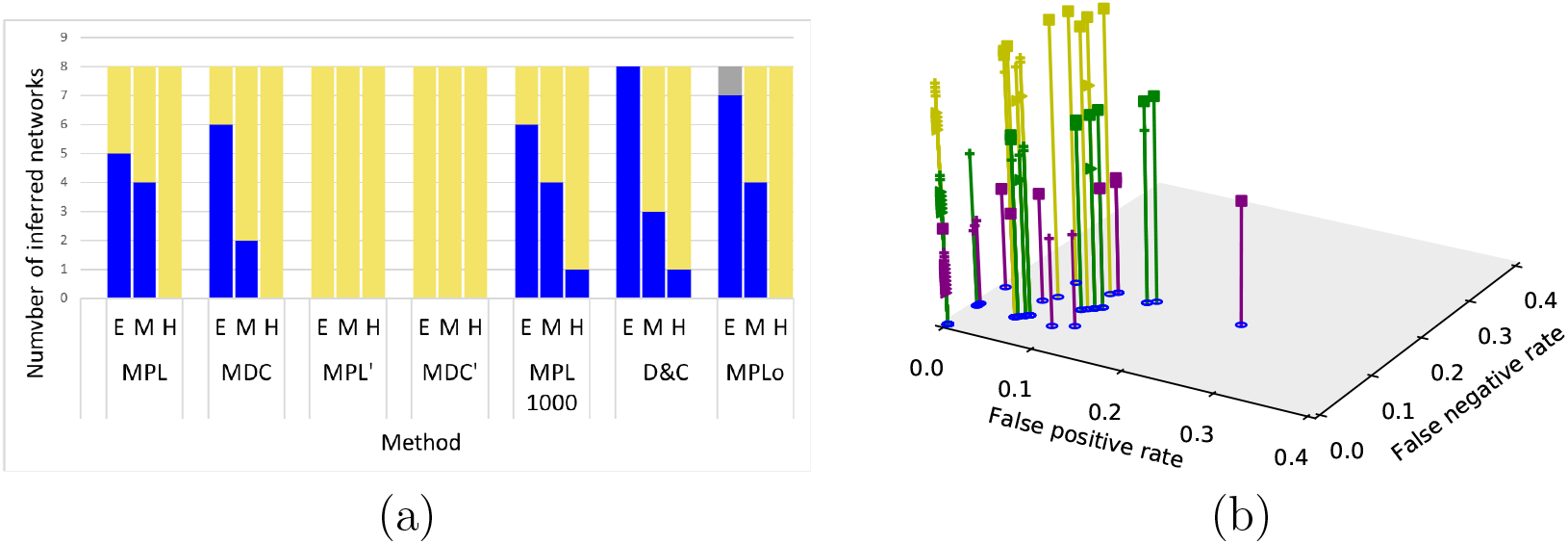
Accuracy of inferred networks. (a) Blue corresponds to the number of inferred networks that are topologically identical to the true networks. Yellow corresponds to the number of inferred networks that share backbone networks of true networks. Gray corresponds to all other cases. (b) Yellow corresponds to MPL. Green corresponds to MDC. Purple corresponds to *D*&*C*. Triangles correspond to easy (E), crosses correspond to medium (M) and squares correspond to Hard (H). The z-axis has no meaning but is used for ease of visualization.

As Figure 2(a) shows, when the true number of reticulations is assumed, both MPL and MDC perform relatively similarly, with MDC outperforming MPL by one of the easy data sets and MPL outperforming MDC by two of the medium data sets. Neither of the two methods obtains the correct network on any of the 8 hard data sets. When the true number of reticulations is not assumed, MPL’ and MDC’s do not infer the correct network on any of the 24 data sets. For all these four methods, when the correct network is not inferred, the distance between the inferred network and true network is *D*(Ψ_*t*_, Ψ_*i*_) = 0. That is, while the methods do not infer the correct network, the inferred networks share an underlying structure with the true ones.

Increasing the number of gene trees in the input to 1,000 results in a slight improvement to MPL, where the true network is inferred on two more data sets. Furthermore, searching the network space directly under the MPL criterion performs almost similarly to a tree-based approach, with the only difference being that the method (MPLo) now infers two more networks correctly and infers a wrong network in the case of the 8 easy networks. It is important, though, to emphasize again that for MPL, MDC, MPL1000, and MPLo, the true number of reticulations is assumed. As the results show, *D*&*C* has the best accuracy where not only does it always either infer the correct network or a network that shares an underlying structure with the true one, but it also does so without *a priori* knowledge of the true or maximum number of reticulations.

To better understand the behavior of these methods, we inspected the FPR and FNR of the methods. As Figure 2(b) shows, on average, the FPR and FNR of MPL are 15.2% and 11.1% higher than those of *D*&*C*, respectively, and those of MDC are 52.1% and 28.2% higher than those of *D*&*C* respectively.

We show the running times of tree-based network inference under both MPL and MDC in Figure 3. The figure shows that almost all data sets are analyzed within 16 hours by MPL when the true number of reticulations is assumed, but that time increases to about 40 hours for some data sets when the search is allowed to explore networks up to 5 reticulations. Inference based on the MDC criterion, on the other hand, takes much longer in some cases, and as the figure shows, the number of reticulations itself is not the only determinant of the running time. The placement of the reticulations in the network is a major factor of the complexity, a result similar to that shown in the case of computing the likelihood of networks [29, 5].

**Figure 3:**
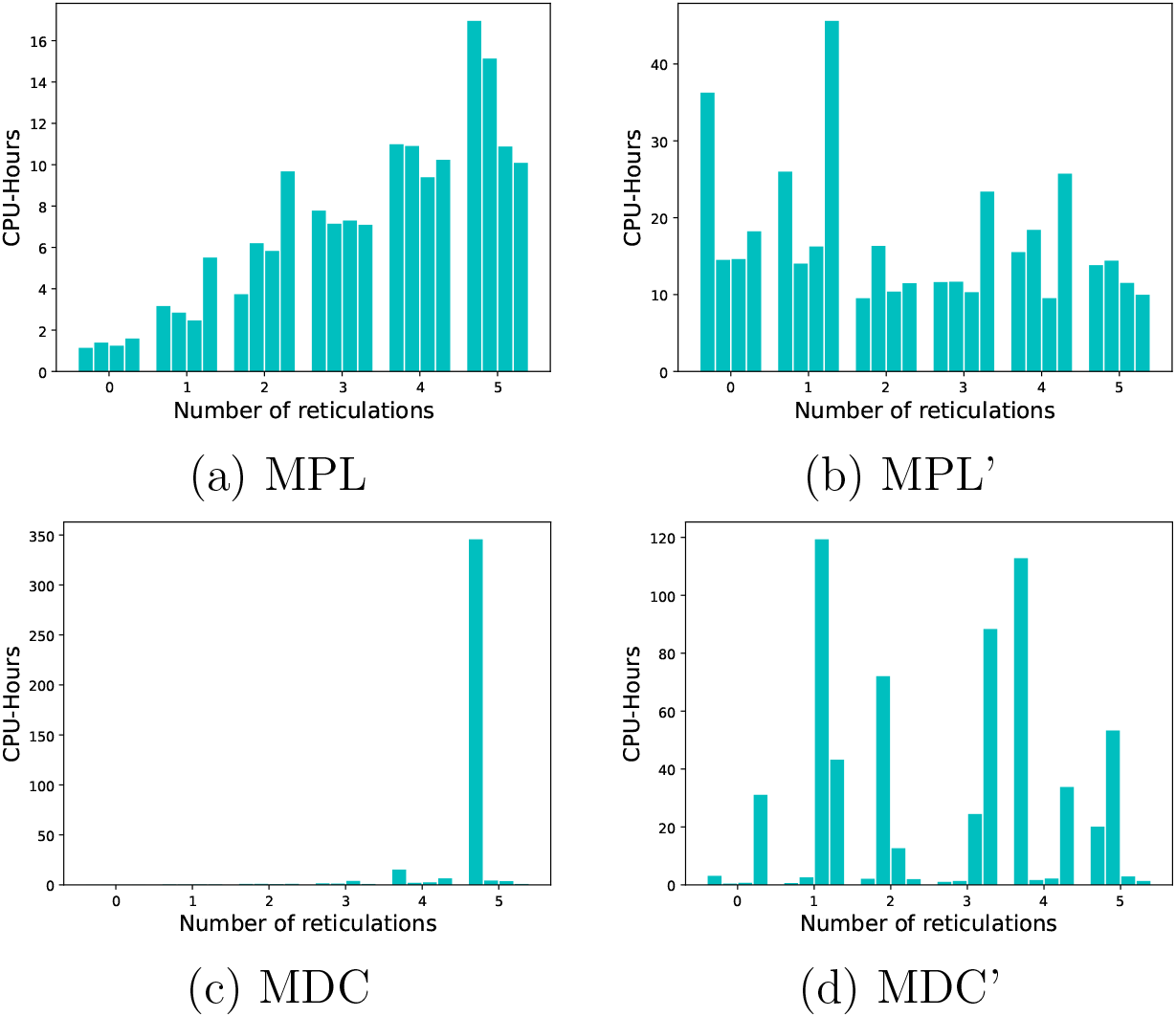
Running times of tree-based network inference based on the MPL and MDC criteria. Each subfigure shows the running times on 24 networks for the respective method. Each bar represents the CPU hours taken to infer this network. The x-axis is the number of reticulations, where each number of reticulations has 4 networks with that number.

On average, when compared with 1636.8 CPU hours spent by *D*&*C*, the tree-augmentation methods only spend 7.0, 16.4, 17.0, 26.4 and 3.8 hours for MPL, MDC, MPL’, MDC’ and MPL1000, respectively. The computational bottleneck of *D*&*C* comes from the substantial time that it takes MCMC-SEQ [16] to infer each of the 680 3-taxon subnetworks (though this can be easily parallelizable, as the inferences of subnetworks are done independently).

### 4.3 Towards combining the strengths of tree-based and *D*&*C* inference

As shown above, tree-based inference is much faster than inference by *D*&*C*, whereas the latter produces more accurate results. As we mentioned, the majority of the running time of *D*&*C* comes from the costly step of inferring the 3-taxon subnetworks using the expensive Bayesian approach of [16]. The question that we set out to explore here is: Can the running time of *D*&*C* be improved by utilizing tree-based inference of the trinets? We limit our attention in this study to one part of this question, namely, how does tree-based inference perform in terms of inferring the 3-taxon subnetwork topologies (when using *a priori* knowledge of the true number of reticulation)? To explore this question, we considered each 3-taxon subset of the 17 taxa in each data set and inferred a network on it (with the true number of reticulation) using tree-based inference (minimizing deep coalescence) starting from all three possible topologies. Figure 4 shows the accuracy and running times of this approach. As the figure shows, on average, the running time of 3-taxon subnetwork inference is reduced from 1636.8 to 1.4 CPU hours with a loss of only about 7% in accuracy when considering identical subnetworks and about 1% when considering backbone networks. This is a massive improvement in the running times with hardly any sacrifice in the topological accuracy. However, there are two caveats here. First, the true number of reticulations is assumed in this case (whereas that number is not assumed in [26]). Second, the merger step of the *D*&*C* method of [26] assumes knowledge of the divergence times of the nodes in the 3-taxon subnetworks. A promising direction that emerges from these results is that combining tree-based inference with the Bayesian method of [16] could potentially provide an accurate and fast approach to inferring the subnetworks and, consequently, improving the running of *D*&*C* without sacrificing its accuracy.

**Figure 4:**
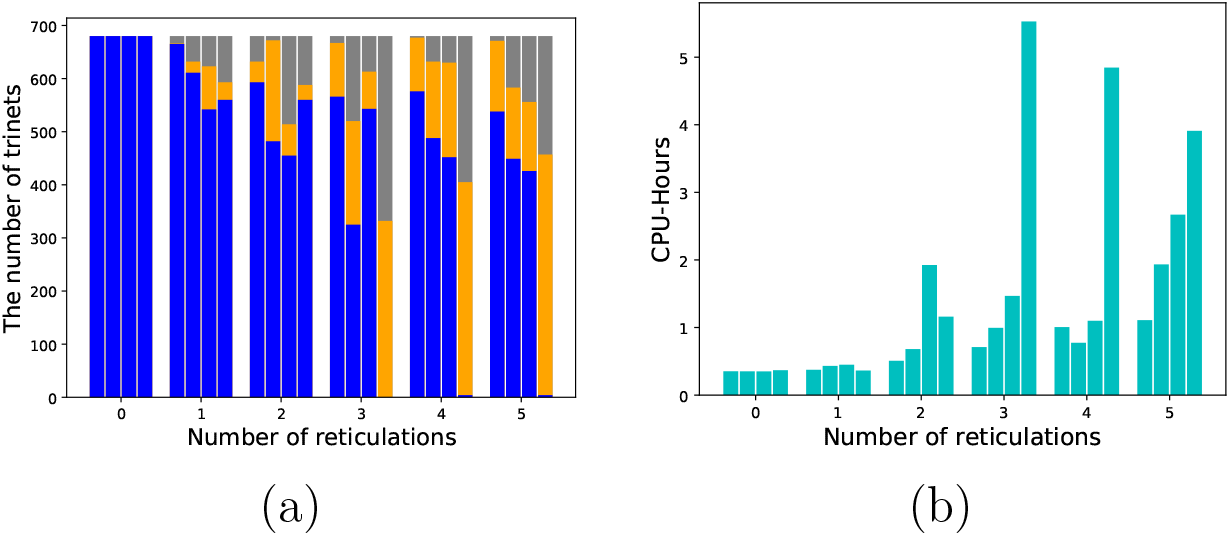
The accuracy and running time of tree-based inference of 3-taxon subnetworks. (a) Accuracy of the inferred subnetworks. Each bar represents the number of subnetworks in each network that are identical (blue), inside (orange), others (grey) to the true subnetworks. (b) Running time for inferring subnetworks. Each bar represents the CPU hours spent to infer all subnetworks for each network. The bars are arranged in a way that they correspond in a 1-1 manner to Figure 3 in [26].

### 4.4 Analysis of empirical data set

We reanalyzed an empirical data set of rainbow skinks [1]. Selecting 11 taxa and 22 individuals, we inferred 100 gene trees from aligned sequences of 100 loci using IQ-TREE. We inferred a start species tree from gene trees using ASTRAL-III, which is shown in Figure 5(a). We then rooted the gene trees and start species trees at the *Lampropholis guichenoti* as an outgroup, and deleted it. We then ran **LocalSearchAugment** under the pseudo-likelihood criterion on the data and set the maximum number of reticulations at 1, in order to compare with the results reported in [26]. The method took 6 minutes to obtain the network shown in Figure 5(b), while *D*&*C* took 3670 CPU-hours. While the start species tree agrees with the inferred species tree reported in Figure 2 in [1], the tree-based network is different from the network inferred by *D*&*C* and reported in Figure 5 in [26], once again, demonstrating the efficiency of tree-based inference, but its limitations in terms of accuracy when run on large data sets.

**Figure 5:**
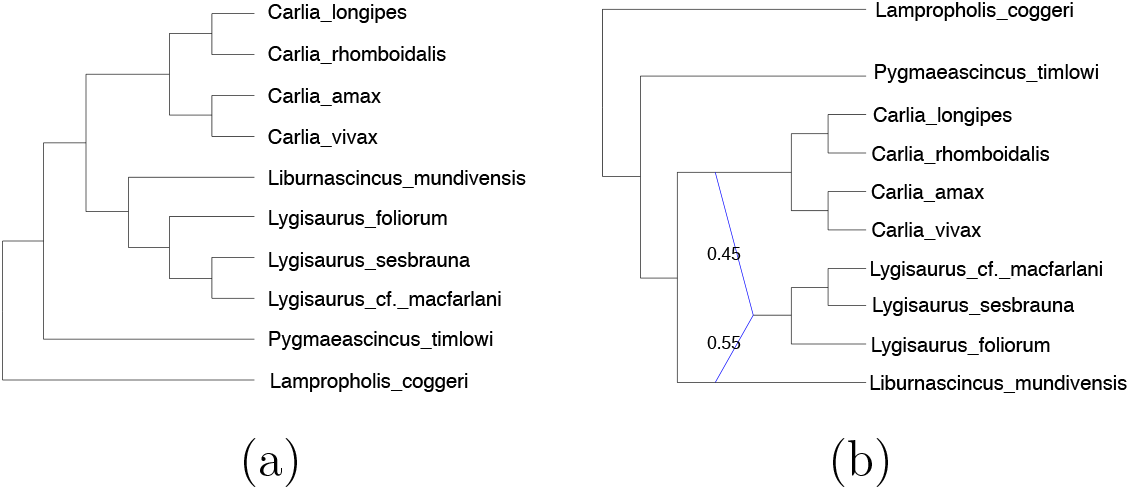
The inferred species tree and network of the empirical data set. (a) The ASTRAL species tree from IQTREE gene trees. (b) The inferred network with 1 reticulation using maximum pseudo-likelihood relying on the backbone tree.

## 5 Conclusions and future work

In this paper, we set out to study the performance of tree-based inference of phylogenetic networks, as this approach would be promising for large-scale phylogenetic network inference provided it has good accuracy. While we find the method to be much faster than a recently introduced divide-and-conquer approach, its accuracy is inferior to the latter. However, the approach is accurate for inference of small-scale networks, which could prove to be valuable for speeding up the divide-and-conquer approach while maintaining its accuracy. For example, the topologies of the subnetworks could be inferred using tree-based inference, and then the Bayesian method of [16] is run to only estimate the divergence times, rather than estimating the topologies as well. We identify this as an important direction for future research.

Another important open question whose answer would have practical implications on searching the network space: is an optimal phylogenetic network with *k* + 1 reticulations (under some optimality criterion) obtainable by adding a reticulation event to an optimal network with *k* reticulations? While the answer to this question could be negative for all optimality criteria (likelihood, pseudo-likelihood, MDC, etc.), the answer could be positive for certain classes of phylogenetic networks.

